# Predicting *N*-strain coexistence from co-colonization interactions: epidemiology meets ecology and the replicator equation

**DOI:** 10.1101/722587

**Authors:** Sten Madec, Erida Gjini

## Abstract

Multi-type spreading processes are ubiquitous in ecology, epidemiology and social systems, but remain hard to model mathematically and to understand on a fundamental level. Here, we describe and study a multi-type *susceptible-infected-susceptible* (SIS) model that allows for up to two co-infections of a host. Fitness differences between *N* infectious agents are mediated through altered susceptibilities to secondary infections that depend on colonizer- co-colonizer interactions. By assuming small differences between such pairwise traits (and other infection parameters equal), we derive a model reduction framework using separation of timescales. This ‘quasi-neutrality’ in strain space yields a fast timescale where all types behave as neutral, and a slow timescale where non-neutral dynamics take place. On the slow timescale, *N* equations govern strain frequencies and accurately approximate the dynamics of the full system with *O*(*N*^2^) variables. We show that this model reduction coincides with a special case of the replicator equation, which, in our system, emerges in terms of the pairwise invasion fitnesses among strains. This framework allows to build the multi-type community dynamics bottom-up from only pairwise outcomes between constituent members. We find that mean fitness of the multi-strain system, changing with individual frequencies, acts equally upon each type, and is a key indicator of system resistance to invasion. Besides efficient computation and complexity reduction, these results open new perspectives into high-dimensional community ecology, detection of species interactions, and evolution of biodiversity, with applications to other multi-type biological contests. By uncovering the link between an epidemiological system and the replicator equation, we also show our co-infection model relates to Fisher’s fundamental theorem and to conservative Lotka-Volterra systems.

## Introduction

Many factors have been shown to be important for ecological biodiversity. These include inter- vs. intra-species interactions (Tilman, 1987), population size (Taylor et al., 2004), number and functional links with resources (Armstrong and McGehee, 1976), and movement in space (Nowak and May, 1992). Diversity of an ecosystem is closely related to its stability, resilience to perturbations and productivity (Hooper et al., 2005). Co-colonization processes appear in many diverse ecological communities, from plant and marine ecosystems to infectious diseases: two species encountering and coexisting together in a unit of space or resource. Interactions between resident and co-colonizer entities upon encounter may exhibit asymmetries, randomness, and special structures, and may range from facilitation to competition. Although special cases for low dimensionality seem to be tractable analytically, the entangled network that arises between *N* interacting entities in co-colonization, and its consequences for multi-type dynamics over short and long time scales remain elusive. Importantly, the question of how the net behavior of a collective of types arises from pairwise outcomes in co-colonization constitutes a nontrivial analytical challenge.

In infectious diseases characterized by polymorphic agents, such as influenza, dengue, malaria, and pneumococcus, the mechanisms allowing many different co-circulating pathogen types to be maintained have long been considered (Gog and Grenfell, 2002; Gupta and Anderson, 1999; Cobey and Lipsitch, 2012a). Typically, strain-specific and cross-immunity have been studied using SIR frameworks, as a main structuring force of such polymorphic systems (Kucharski et al., 2016). In contrast, the influence of interactions through co-colonization (or co-infection, implying simultaneous carriage of two strains), which reaches for example 10-20% prevalence in asymptomatic carriage in pneumococcus (Valente et al., 2012), has received less mathematical attention on the SIS modeling spectrum (Adler and Brunet, 1991; Lipsitch, 1997; Gjini et al., 2016). In pneumococcal bacteria, displaying more than 90 serotypes (Park et al., 2007), carriage of one serotype has been shown to alter, mostly reduce, the acquisition rate of a second serotype (Auranen et al., 2010; Lipsitch et al., 2012). Despite competition, much diversity at the serotype and genotype level persists over long time, even in the face of vaccines (Weinberger et al., 2011). Understanding how co-colonization interactions may promote coexistence patterns within and between species, in pneumococcus (Bogaert et al., 2011; Dunne et al., 2013; Shrestha et al., 2013)) or other pathogen systems (Cohen et al., 2008; Abdullah et al., 2017) remains a challenge.

To meet this challenge, mathematical frameworks need to integrate diversity and inter-dependence in such multi-type systems with ecological principles and simpler formulations. Unfortunately however, mechanistic insights for high-dimensionality are not always available. In pneumoccocus, previous epidemiological studies, when including co-colonization, have either adopted low system size for analytical results (*N* = 2) (Lipsitch, 1997; Gjini et al., 2016), focusing on vaccination outcomes, or combined niche and neutral mechanisms exclusively through simulations for larger number of strains (large *N*) (Cobey and Lipsitch, 2012a; Bottomley et al., 2013; Nurhonen et al., 2013). In other systems, advances for *N* = 3 and special parameter combinations, have only recently been made (Pinotti et al., 2019).

To advance our understanding and predictability of multi-type systems, there is a need to overcome both these limitations, increasing our analytical power over a larger and more realistic system size. For example, *N* ≈ 30 is a typical number of co-circulating pneumococcus serotypes in any given setting (Lipsitch et al., 2012), but comprehensively, the quadratic explosion of number of serotype pairs with *N* makes it very hard to predict and mechanistically understand epidemiological variables describing co-colonization.

Limitations in modeling interacting disease systems persist even in more general theoretical contagion studies of SIS (Hébert-Dufresne and Althouse, 2015; Chen et al., 2017) and SIR type (Sanz et al., 2014; Miller, 2013), which have recognized the rich behavior of such systems, especially when cooperative interactions between entities are at play. These studies have also considered very low system size (*N* = 2), and either only cooperative or only competitive interactions, highlighting more critical transitions in contact networks (Hébert-Dufresne and Althouse, 2015; Miller, 2013). Joint analyses of cooperation and competition under the same framework, for an arbitrary number of interacting entities, that avoid combinatorial explosion of complexity, remain undeveloped.

In this study, we provide a fundamental advance on this challenge. We describe and analyze a very general system of multi-type interactions that arise in SIS epidemiological dynamics with co-infection (co-colonization). We show that by assuming small differences in altered susceptibilities to coinfection between *N* types, an explicit reduced system emerges via a timescale separation. This enables to express the total dynamics as a fast (neutral) plus a slow (non-neutral) component, the latter related to variation in co-colonization interactions. Multi-strain collective dynamics can be very complex, but we find that they are explicitly modulated by global parameters such as overall transmission intensity *R*_0_, and mean interaction coefficient between strains, *k*. We derive a closed analytic solution for strain frequencies over long time-scales in a changing mean fitness landscape. We further show that the emergent equation from this *N*-dimensional model reduction corresponds to a version of the classical replicator equation from game theory (Taylor and Jonker, 1978; Weibull, 1997; Hofbauer and Sigmund, 2003), in our case, appearing in terms of pairwise invasion fitnesses between strains.

Although the model is treated in an epidemiological spirit, parallels and conceptual analogies with other systems can be easily drawn, where multiple types (species/strains/entities) in a homogeneous mixing scenario compete for free and singly-occupied niches via generic colonizer-cocolonizer interactions. In the paper, we will use the terms co-infection and co-colonization interchangeably, referring always to an avirulent scenario, and thus reinforcing the potential application of the framework both in epidemiology and ecology.

## The modeling framework

### N-strain SIS model with co-colonization

We consider a multi-type infectious agent, transmitted via direct contact, following susceptible-infected-susceptible (SIS) transmission dynamics, with the possibility of co-infection. The number of potentially co-circulating strains is *N*. With a set of ordinary differential equations, we describe the proportion of hosts in several compartments: susceptibles, *S*, hosts colonized by one type *I*_*i*_, and co-colonized hosts *I*_*ij*_, with two types of each combination, independent of the order of their acquisition. We have:

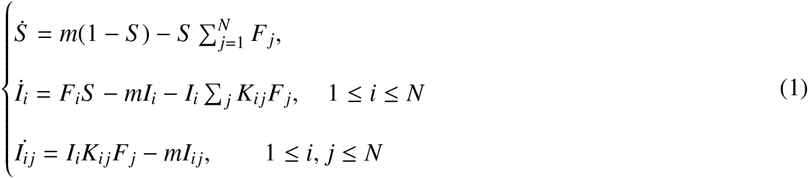

where 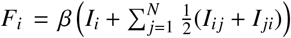 gives the force of infection of strain *i*. We assume that hosts colonized with a mixture of two subtypes *i* and *j*, *I*_*ij*_ transmit either with equal probability. For more information and interpretation of parameters see Table S1. Notice that *S* = 1 − ∑ (*I*_*i*_ + *I*_*ij*_), thus the dimension of the system is indeed *N* + *N*^2^. In practice, *I*_*ij*_ and *I*_*ji*_ cannot be distinguished so the dimension is effectively *N* + *N*(*N* − 1)/2.

We study pathogen diversity only in relation to how strains interact with each other upon co-colonization (*K_ij_*), assuming equivalence in transmission *β* and clearance rate *γ*. The coefficients *K*_*ij*_, denote factors of altered susceptibilities to secondary infection when a host is already colonized, and can be above or below 1, indicating facilitation or competition between colonizer and co-colonizer strain. In the above notation *m* = *γ* + *r*, encapsulating both clearance rate *γ* of colonization episodes and recruitment rate of susceptible hosts *r*, equal to the mortality rate from all compartments (*r* = *d*). For a model summary diagram see Fig.1a-b. Since we model *N* closely-related strains, we can write each co-colonization coefficient as: *K*_*ij*_ = *k* + *εα*_*ij*_, where 0 ≤ *ε* ≪ 1. This will form the basis of our model reduction framework, and system decomposition into smaller sub-systems.

**Figure 1:**
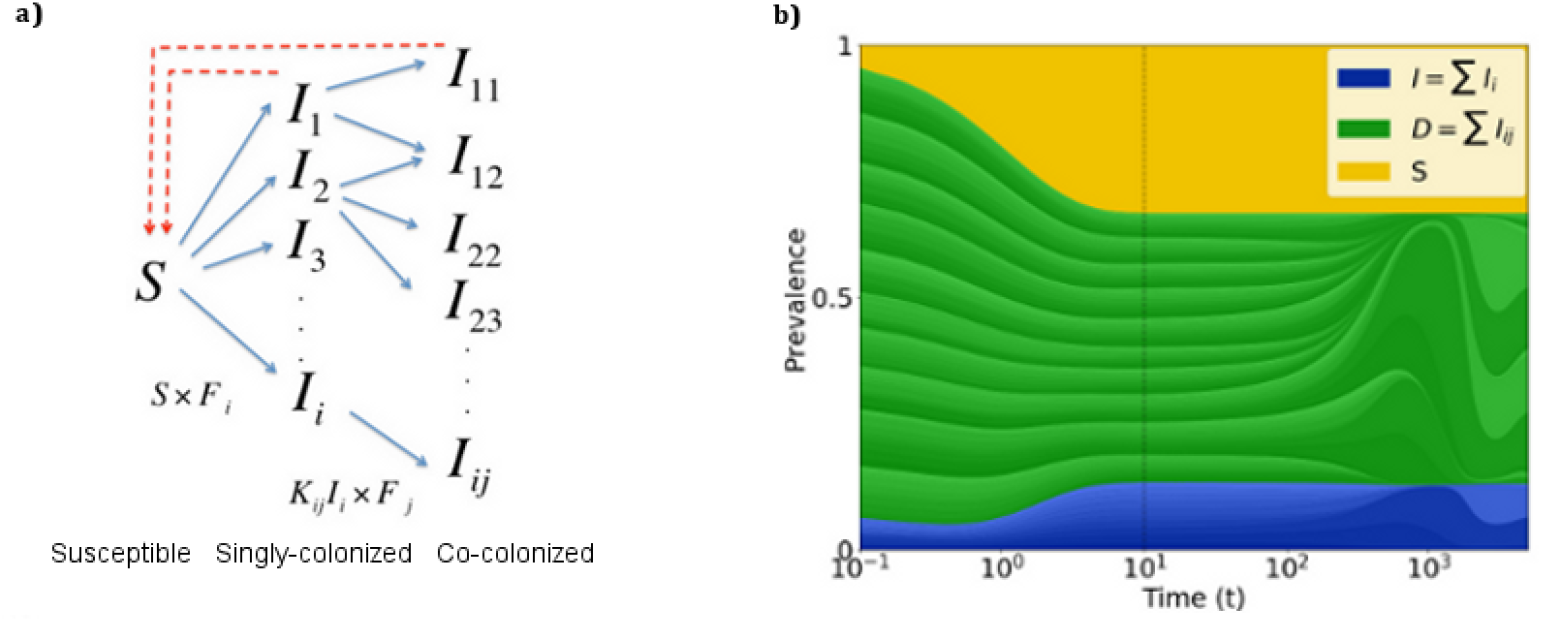
Model summary diagram. **a. Co-colonization model structure.** Hosts move from susceptible to singly-colonized state, and from singly-colonized to co-colonized state. Clearance happens at equal rate for single and co-colonization. Co-colonization rate by strain *j* of singly-colonized hosts with *i* is altered by a factor *K*_*ij*_ relative to uncolonized hosts. **b. Complex epidemiological dynamics can be represented in two interrelated timescales.** Assuming that pairwise interaction coefficients in co-colonization can be represented as: *K*_*ij*_ = *k* + *εα*_*ij*_, the global compartmental dynamics can be decomposed in a fast and slow component. On the fast time-scale (*o*(1/*ε*)), strains follow neutral dynamics, driven by mean-field parameters, where total prevalence of susceptibles *S*, singly-infected hosts, *I* and dually-infected hosts, *D*, are conserved. On a slow time-scale, *εt*, complex non-neutral dynamics between strains takes place, depicted here by the constituent variations within the blue and green. These non-neutral dynamics are explicitly derived in this paper, and yield an explicit equation for strain frequency dynamics.

## Results

### Re-writing the system in aggregated form

In order to derive a reduced model for such a general *N* strain system, displaying complicated and high-dimensional dynamics (Fig. 1b), we use the following aggregation of variables:

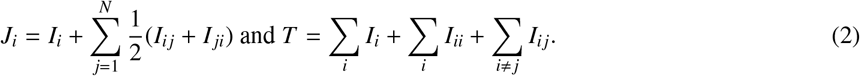

for the prevalence of strain *i* in the population, and the total prevalence of all strains, respectively. Total prevalence satisfies *T* = ∑_*i*_ *J*_*i*_ and the forces of infection are: *F*_*i*_ = *βJ*_*i*_. With these notations, the system (1) can be rewritten as:

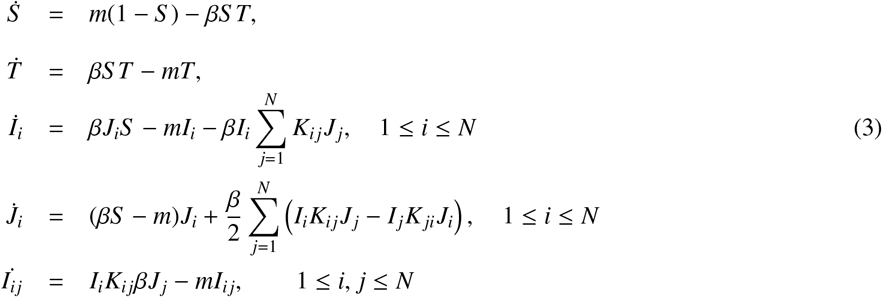

This system of 2 + *N* + *N*^2^ equations, now displays a convenient structure, that we exploit for our analysis: i) First we describe the block of 2 equations (*S*, *T*) that do not depend on *K*_*ij*_. ii) Next, we study the block of 2*N* equations (*I*_*i*_, *J_i_*), which is the most complicated. iii) Lastly, we deal with the block of the *N*^2^ equations of *I*_*ij*_, which is simple once the dynamics of *I*_*i*_ and *J*_*i*_ are known (see Supplementary material 2 for details). Clearly, if the basic reproduction number 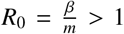, as typical in SIS models, then there is an endemic equilibrium, whereby 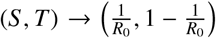 (Dietz, 1993). By reducing the system to the invariant manifold 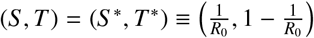, we obtain a simpler system for the second block of equations. Once it is known that 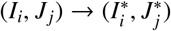, then the *N*^2^ equations for co-colonization compartments imply 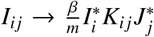. Thus, once the dynamics of the second set of equations are explicit, so are the dynamics of co-colonization variables, and ultimately of the entire system.

### Slow-fast decomposition of the system

We use a similar trick as in (Gjini and Madec, 2017). We assume the *N* strains are closely-related, thus each co-colonization coefficient can be rewritten as: *K*_*ij*_ = *k* + *εα*_*ij*_, where 0 ≤ *ε* ≪ 1, and *k* a suitable reference. Replacing these in (3), re-arranging, and analyzing the cases of *ε* = 0 and *ε* > 0 but small, we obtain the slow-fast representation of the system (See Materials and Methods, and Supplementary Material 2 for details).

#### Neutral system

If *ε* = 0, then we obtain the *Neutral model*, where all strain co-colonization coefficients are equal to *k* and strains behave as equivalent. We find that on the fast time-scale *o*(1/*ε*), the key dynamics are given by these two quantities:

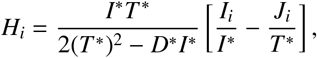

and

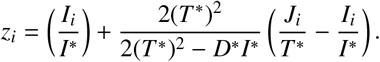

which obey: 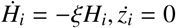. The quantity *H*_*i*_ measures the difference between the part occupied by strain *i* in single colonization versus the part of strain *i* in total carriage. Thus, *H*_*i*_ → 0 means that, on the fast timescale, the proportion of strain *i* in single colonization 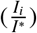, tends to equalize the proportion of strain *i* in overall colonization 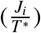. This implies it also tends to be equal to the proportion occupied by strain *i* in co-colonization: 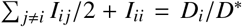. Because *H*_*i*_ = 0 is the only equilibrium, we can infer that during fast dynamics *z*_*i*_ tends to the limit:

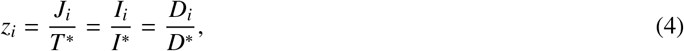

whereby *z*_*i*_ becomes the variable in the system that exactly describes exactly the frequency of strain *i* in the host population, with ∑ *z*_*i*_ = 1.

#### The slow dynamics: strain frequencies

Analyses of the system for *ε* > 0, reveal that on the slow timescale *τ* = *εt*, strain frequencies, *z_i_*, obey explicit dynamics (See Supplementary material 3). These dynamics are given by the *N*-dimensional system:

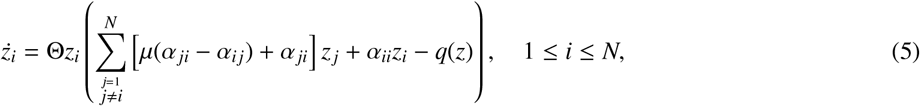

where the constants Θ, *μ* > 0 are explicit functions of the global steady state (*T*\*, *I*\*, *D*\*) of the neutral model:

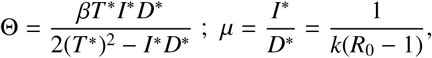

and the term *q*(*z*) is a quadratic term given by:

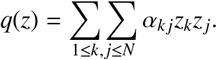

The above expression encapsulates how the ultimate competitive hierarchies between *N* co-colonizing strains, circulating in the same host population, are driven by the magnitude of asymmetries in within-vs. between-strain interactions (the *α*’s), as well as by mean-field global quantities, such as *R*_0_ and *k*, affecting Θ and *μ*. Thus, the strain selection occurring in the slow time scale, on the basis of small differences in co-colonization interactions, becomes entirely explicit.

From (5), we can notice that these equations resemble conservative Lotka-Volterra dynamics (Lotka, 1926; Volterra, 1926), but with an extra term *q*(*z*), which changes during time. This term *q*(*z*) represents the evolving impact of all the strains on their ‘common environment’, which in turn modifies their own fitness landscape. A more explicit way to interpret *q*(*z*) is in terms of relative change in ‘effective’ mean interaction coefficient between all extant types in the system, which if negative, indicates a global trend toward more competition in co-colonization, and if positive, indicates a global trend toward more facilitation. Formally one would write this mean trait dynamics as 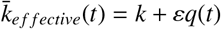.

### Link with pairwise invasion fitnesses 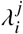 and the replicator equation

Next, we uncover an equivalent very useful representation of the model reduction by using the notion of pairwise invasion fitness (Metz et al., 1992; Geritz et al., 1998; Meszéna et al., 2005). Let 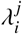 be the exponential growth rate of strain *i* evaluated when introduced at the trivial endemic equilibrium of the strain *j* alone. If the fitness 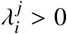, strain *i* will invade *j*, and viceversa. By considering the rate of growth of strain *i* in an endemic equilibrium set by *j* (*dz*_*i*_/*dt* (5) in the special case where all *z*_*k*_ = 0 for *k* ≠ *j*), we find the exact formulation of pairwise invasion fitness, in our model, is given by:

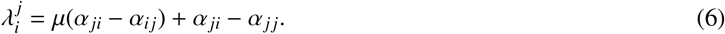

We can thus recast the strain frequency equations (system (5)) using only these fitness notations (for details see Supplementary Material 3):

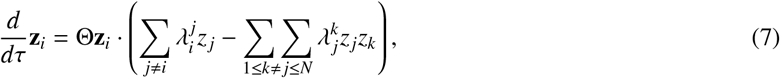

which can be made even more compact by denoting the pairwise invasion fitness matrix 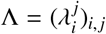, and using vector notation:

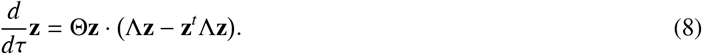

Ultimately, it is this matrix Λ that defines all ‘edges’ of the rescaled interaction network between *N* strains. For *N* = 2 and 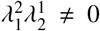, similar to the classical competitive Lotka-Volterra model, in our model, as previously shown (Gjini and Madec, 2017), there are only four possible outcomes between 2 strains (edge linking 1 and 2): **i)** 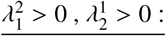 stable coexistence of 1 and 2; **ii)** 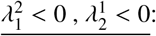 bistability of 1-only and 2-only; **iii)** 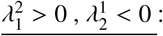 1-only competitive exclusion; **iv)** 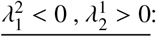 2-only competitive exclusion. Knowing all pairwise invasion fit-nesses between each couple of strains, via expression (8) we can reconstitute the ultimate dynamics of the full system with *N* types and co-colonization. Seen more closely, this model reduction in strain frequency space, corresponds to an instance of the classical replicator equation for multiplayer games, long studied in evolutionary game theory (Taylor and Jonker, 1978; Weibull, 1997; Hofbauer and Sigmund, 2003). Remarkably, however, in contrast to assuming it heuristically *a priori*, here we have derived it from basic aggregation and timescale principles in an epidemiological context. Moreover, the pairwise traits in the emergent expression are indeed special traits, denoting pairwise invasion fitnesses between co-colonizing strains; invasion fitness being a central quantity in adaptive dynamics used to predict trait evolution(Metz et al., 1992; Geritz et al., 1998; Meszéna et al., 2005).

The entire dynamics of the *N*-strain ‘game’ can now be recapitulated based only on knowledge of the pairwise invasion network between each two ‘players’. By having heterogeneous interactions across strain space, and receiving heterogeneous interactions from other strains in the network, each strain, competing for susceptibles and singly-colonized hosts, “perceives” its own unique environment, creates its own niche, which ultimately enables its persistence or extinction. However, the global fitness of each strain in such coupled multi-player competition, depends not only on its own individual fitness and frequency at any time, but also on the fitnesses and frequencies of all other strains. This feature can lead to a multitude of outcomes, as already recognized in replicator equation studies (Nowak and Sigmund, 2004; Cressman and Tao, 2014).

For illustration, in Figure 2 and in Supplementary Movie S1, we provide an example of the modeling framework and the coexistence dynamics that arise among a number of strains (here *N* = 6) for an arbitrary co-colonization interaction matrix *K*. Another combination of parameters, leading to a limit cycle for *N* = 6, is illustrated in Supplementary Figure S1 and Supplementary Movie S2. These examples demonstrate that even weak asymmetries between apparently similar types, in altered susceptibilites to co-colonization, have the potential to generate rich and hierarchical collective behavior over long time.

**Figure 2:**
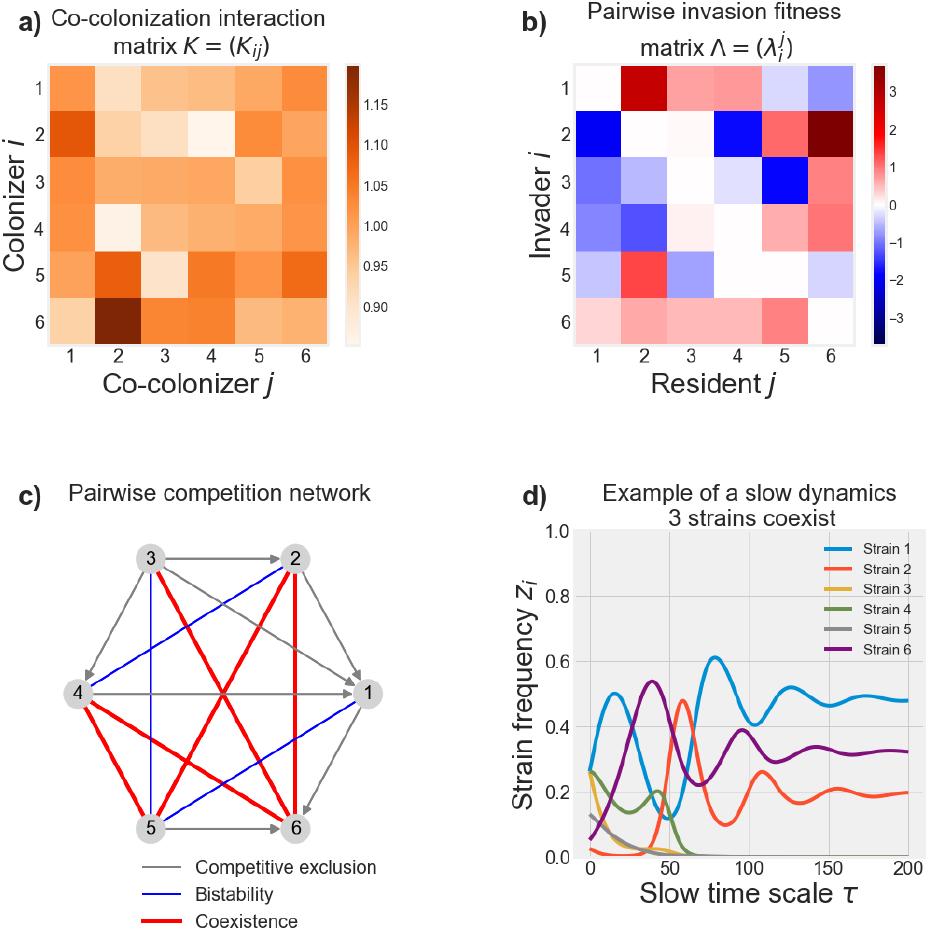
Example dynamics of our model for *N* = 6. **a)** The matrix of interaction coefficients in co-colonization, generated randomly, with mean *k* = 1, and standard deviation *ε* = 0.1. **b)** The corresponding pairwise invasion fitness matrix for assumed *R*_0_ = 2. **c)** The network where each edge displays the outcome of pairwise competition between any couple of strains, where the direction of grey edges denotes the winner in competitive exclusion. **d)** Slow epidemiological dynamics resulting from these interactions. A dynamic display of the trajectory is shown in Supplementary Movie S1.

Note that in Equations (5) and (8), the constant Θ is the rate that sets the tempo of multi-type “motion” on the slow timescale towards an equilibrium. The pre-factor Θ depends specifically on the absolute transmission rate of the pathogen *β*, which calibrates the slow dynamics, but also on mean traits *R*_0_ and *k*, via the conserved aggregated quantities *T*\*, *I*\*, *D*\*, where *R*_0_ and *k* are nonlinearly coupled (see Supplementary Figure S2). This shows how global environmental variables (similar to the notions of effective population size in population genetics) affect the speed of non-neutral dynamics between types in the system. The quantity *μ* = *I*\*/*D*\* on the other hand represents the ratio between single colonization and co-colonization prevalence in the overarching neutral system, a critical factor that amplifies the importance of type asymmetry in the slow dynamics due to deviation from neutrality (Gjini and Madec, 2020). A summary of key model quantities in terms of mean field parameters *R*_0_ and *k* is given in Table 1.

**Table 1:** Key model quantities in terms of basic reproduction number *R*_0_ and reference interaction coefficient in co-colonization *k*.

| Symbol | Interpretation | Features |
| --- | --- | --- |
| $R_0$ | Basic reproduction number | $R_0 = \beta/m > 1$ |
| $k$ | Reference interaction coefficient between types in co-colonization (e.g. mean of $K_{ij}$) | $k \begin{cases} < 1 : \text{competition,} \\ > 1 : \text{cooperation} \end{cases}$ |
| $S^*$ | Equilibrium prevalence of susceptibles | $S^* = \frac{1}{R_0}$ |
| $I^*$ | Equilibrium prevalence of singly-colonized hosts | $I^* = \frac{R_0 - 1}{R_0[1 + k(R_0 - 1)]}$ |
| $D^*$ | Equilibrium prevalence of co-colonized hosts | $D^* = \frac{(R_0 - 1)^2 k}{R_0[1 + k(R_0 - 1)]}$ |
| $\mu$ | Ratio between single and co-colonization | $\mu = \frac{I^*}{D^*} = \frac{1}{(R_0 - 1)k}$ |
| $\Theta$ | Rate of slow dynamics for strain frequencies ($z_i(\tau)$) | $\Theta = \beta \left( 1 - \frac{1}{R_0} \right) \left( \frac{\mu}{2(\mu + 1)^2 - \mu} \right)$ |
| $I_{ij}$ | Co-colonization prevalence with $i$ and $j$ | $I_{ij} = kR_0[1 + k(R_0 - 1)]I_i I_j$ |

### The quadratic term Q and evolution of mean fitness

In the equivalent representation of the slow dynamics equation, (8), the common quadratic term *Q* acting on each strain’s frequency, relates the common changing environment to pairwise invasion fitnesses (Metz et al., 1992; Geritz et al., 1998): 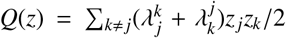. This sum over all extant pairs in the system reflects the mean ‘pairwise invasibility’ of the system as a whole, changing over time with strain frequencies *z*_*j*_(*τ*), *z*_*k*_(*τ*). Upon closer inspection of (8), information on the resilience of a group of strains may be derived from the sign of *Q*: if *Q* > 0 then each existing strain is less competitive within the group, but the overall community is more resistant to invasion by a new outsider strain, and viceversa, if *Q* < 0, then each existing strain grows more within the group, but the overall community would inevitably be also more permeable to invasion by invader strains. This feature of the model, embodies an instance of a classical trade-off between individual and group-selection in evolution (Michod, 2000).

Next, we highlight the meaning of *Q* by considering a few special cases for the invasion fitness matrix between strains (see Supplementary Material 4), illustrated in Figure 3:

i. *<u>Symmetric matrix.</u>* In fact, a symmetric 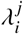 structure between strains in mutual invasion leads to a general feature of the dynamics whereby *Q* always increases over time (Fig. 3a). This case of our *N*-dimensional system 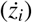, namely the replicator equation for doubly symmetric games, is formally equivalent to the continuous time model of natural selection at a single (diploid) locus with *N* alleles, known as Fisher’s fundamental theorem of natural selection (Fisher, 1958; Price, 1972; Edwards, 1994). In this case, it can be shown that the population mean fitness increases over time, with the rate of change in mean fitness equal to the trait variance at any point (see S4 for full verification of this feature also in our version of this model). In our modeling context, where *λ*_*ij*_ denote pairwise invasion fitness between any two strains, the increase in mean fitness during selective dynamics among *N* strains, implies that when pairwise invasion ‘games’ are symmetric, the system becomes more resistant to invasion by outsider strains over time.
ii. *<u>Invader-driven invasion.</u>* In this case, columns of Λ are equal, meaning it’s differences in ‘attack rates’ (invasiveness) of types that are defining their hierarchical dynamics (Fig. 3b). Mean fitness *Q* again evolves over slow time, reflecting the selection occurring in the multi-type system, and again tends to increase toward positive values, suggesting coexistence is more likely, although in special cases competitive exclusion may occur.
iii. *<u>Resident-driven invasion.</u>* In this case, multi-strain dynamics are driven by variation in ‘defense’ or invasability (rows of Λ are equal), and the principle of competitive exclusion (with possible multi-stability) applies more often, whereby the weakest strains are excluded, and only the best ‘defender’ of its territory (equilibrium when alone) survives. Competitive exclusion obviously implies *Q* should tend to 0, verified in Fig. 3c. In exceptional cases coexistence may also be possible in this case (see S4 for details).
iv. *<u>Antisymmetric matrix.</u>* This is the case when 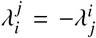 and the propensity for complex coexistence dynamics between strains is very high (Fig. 3d). *Q* is exactly zero in this case, corresponding to zero-sum-games in evolutionary game theory (Hofbauer and Sigmund, 2003). Like in the classical prey-predator Lotka-Volterra system, there exists a unique center surrounded by a family of cycles Chawanya and Tokita (2002). This type of oscillatory coexistence is structurally unstable.
v. *<u>Almost-antisymmetric.</u>* In this case, a small perturbation of the pure anti-symmetric structure in mutual invasion fitnesses disrupts the center leading to a stable or an unstable node. This gives rise to positive and periodic *Q*, where limit-cycles, heteroclinic cycles or chaos are more likely for multi-strain coexistence.
vi. *<u>Random mutual invasion.</u>* In this case, which is the most general case, captured by our framework, the dynamics of *Q*(*z*) can be arbitrary, and increase or decrease over the same realization of multi-strain dynamics, thus encapsulating dynamic shifts in ‘environment quality’, and unpredictable emergent dynamics of mean fitness over time (Fig. 3e). When comparing these 6 special cases, we find that over many stochastic realizations of matrices with such canonical structures and initial conditions, the symmetric invasion matrix results in fastest global mean increase in fitness, as opposed to the random case which leads to the slowest increase in *Q* (see Supplementary Figure S3), in support of the strong stabilizing effect of pairwise symmetry for collective community dynamics.

**Figure 3:**
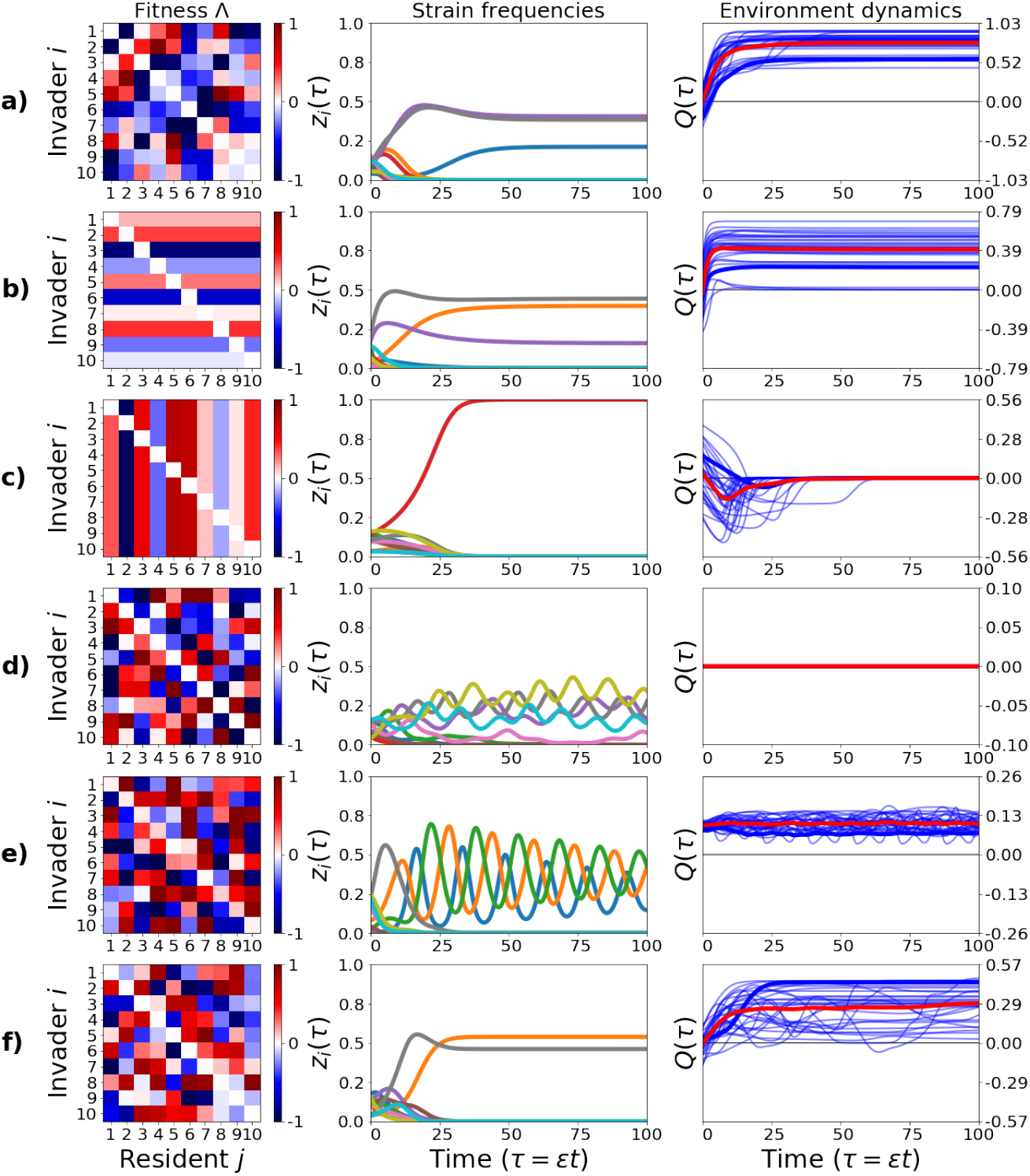
Canonical pairwise invasion structures (Λ) between *N* types and collective dynamics evolution. We generated random 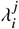 matrices from 6 special cases and simulated multi-type dynamics (*N* = 10) under many realizations of the model ((8)), starting from random initial conditions on the slow manifold, for each case. *Q* is the mean fitness term in the system (the common ‘environment’ for all types) changing differently depending on the pairwise invasion fitness matrix. In the third column, the thin blue lines indicate *Q* evolution for each realization, the thick blue line indicates *Q* evolution for the *z*_*i*_ dynamics shown in the second column, and the thick red line depicts the mean over all 30 realizations. **a) Symmetric matrix.** This corresponds to the same dynamics captured by Fisher’s fundamental theorem. **b) Invader-driven** fitnesses (‘hierarchical attack’). Large potential for coexistence. **c) Resident-driven** fitnesses (‘hierarchical defense’). Large potential for competitive exclusion. **d) Anti-symmetric** invasion fitnesses. *Q* is exactly zero over all time and there is large potential for complex multi-strain behavior. **e) Almostantisymmetric** invasion fitnesses. Maintenance of potential for complex dynamics (e.g. limit cycles) leading to periodicity (but positivity) in *Q*. **f) Random** mutual invasion. Rich model behavior is possible. On average coexistence is more likely, but increases as well as decreases in *Q* over a single realization are possible.

It is worth noting that special cases of *Q* in 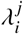 space are more straightforward to analyze and understand than special cases in *K*_*ij*_ trait space, because for each 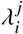 representation (see Eq. (6)), there is an infinite number of possible *K*_*ij*_ in this model, leading to the same pairwise invasion network between strains. In particular, when co-colonization coefficients *K*_*ij*_ display a row-wise or column-wise structure (see (Lipsitch et al., 2012) for such hypotheses in pneumococcus), invoking a strain-specific hierarchy in this trait, for *N* = 2 the principle of competitive exclusion applies, but for general number of strains *N*, such case collapses to the *Q* = 0 case (antisymmetric invasion matrix above) and complex dynamics are expected.

### Going back to the original *N* + *N* (*N* − 1)/2 epidemiological variables

Now let us make the link to the original system with *N* strains, given by the SIS model with co-colonization interactions (1). Assume the *K*_*ij*_ and the global epidemiological parameters are known. The aim here would be to use the model reduction for computational reasons or for analytical insights. Notice that the framing *K*_*ij*_ = *k* + *εα*_*ij*_ is mathematically non-unique, and can be applied with respect to any reference *k*, provided that the resulting *ε* is small. However, one convenient choice is to define *k* as the average of the original interaction matrix entries *K*_*ij*_:

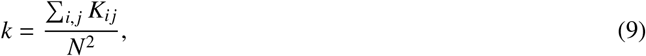

and to define deviation from neutrality, *ε*, as the root mean square distance of the *K*_*ij*_ from the mean interaction coefficient *k*:

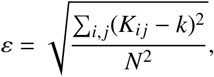

thus representing the standard deviation of the *K*_*ij*_ traits in the pool of *N* available strains. The direction of deviation from neutrality (bias) for the interaction between strain *i* and *j* is then obtained as:

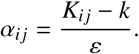

Thus, the matrix *A* = (*α*_*ij*_)_1≤*i*, *j*≤*N*_ is the *normalized interaction matrix*, with 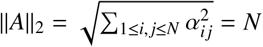. This matrix *A*, and the ratio *μ*, determine the pairwise invasion fitness matrix (Eq.(6)), which contains nearly all the qualitative information about the non-neutral dynamics (Eq.(8)). So far, we have shown that provided the deviation from neutrality, *ε*, is small, the behavior of our approximation (5) describes very well the long term dynamics of the original system (1). To recover the original variables, after solving for strain frequencies *z*_*i*_ on the slow manifold, as the dynamics in Eqs.(1) are well approached by those in Eqs.(5), we can use the relations:

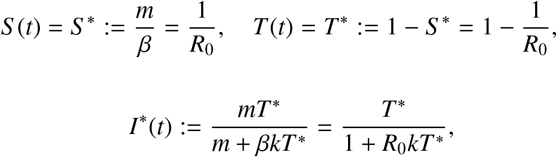

to obtain the total prevalence of uncolonized hosts *S*, total prevalence of colonized hosts *T*, and prevalence of single colonization *I*. The prevalence of co-colonization follows simply from *D*\* = *T*\* − *I*\*. Further, to recover strain-specific single colonization, and co-colonization prevalences in the system, within such a ‘conservation law’, reminiscent of other conservation laws in ecology (Hubbell, 2001), we have:

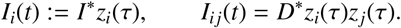

where the slow time scale is *τ* = *εt*, and the strain frequencies *z_i_*(*τ*) verify ∑_*i*_ *z*_*i*_(*τ*) = 1 and follow explicit dynamics (8). The recovery of the epidemiological variables from the replicator equation exposes two special features of the multi-strain dynamics: on one hand, the slow variables *z*_*i*_ (1 ≤ *i* ≤ *N*), describing relative strain frequencies in the host population, tend to be necessarily equal in single and co-colonization (Figure 4a); on the other hand, the prevalence of co-colonization with strains *i* and *j* in this system (Figure 4b), is proportional to the product between single prevalences of *i* and *j* in the population (*I*_*ij*_ ~ *I*_*i*_*I*_*j*_). Notice, that there is an explicit mean-field pre-factor, dependent on *R*_0_ and *k*, determining this constant of proportionality (see Table 1). These two quasi-neutrality principles, are preserved on the slow timescale, independently of strain identities and for all time, and moreover, independently of the complexity of the dynamics.

**Figure 4:**
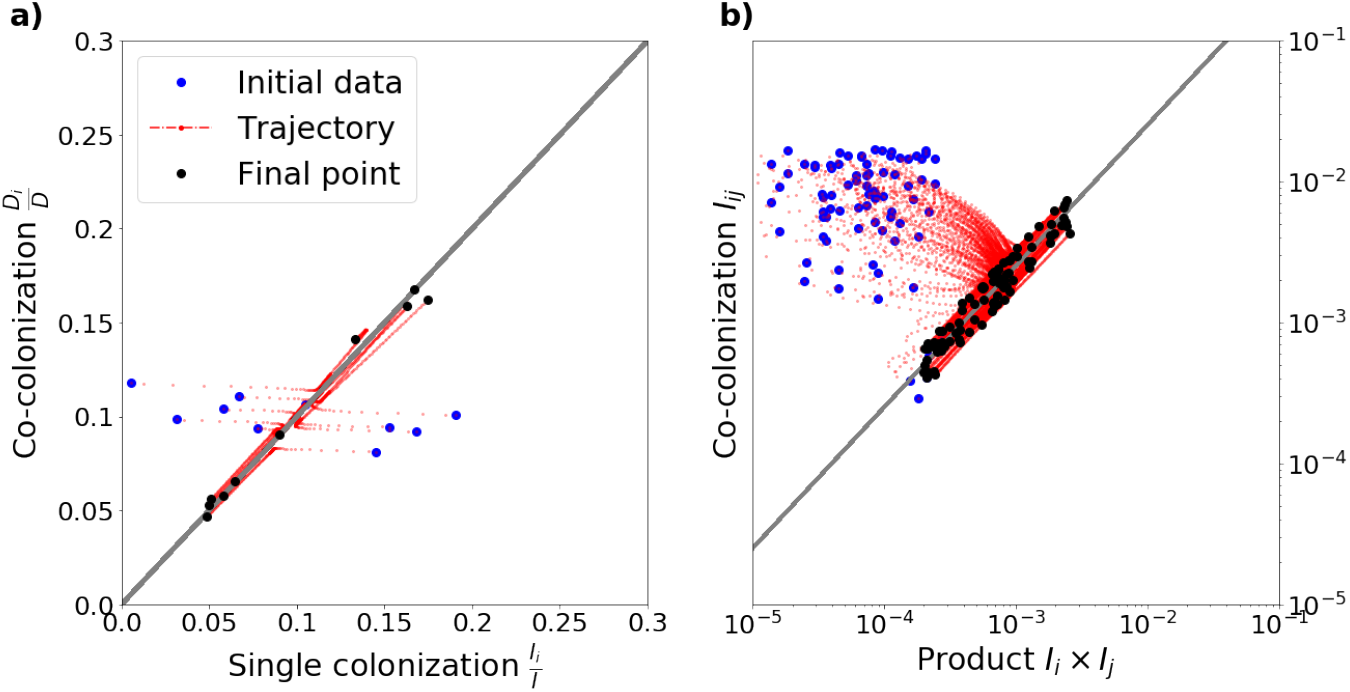
Illustration of invariant principles for strain coexistence on the long timescale. **a)** Strain frequencies tend to be the same in single and co-colonization, for all strains and for all time when on the slow manifold ((4)). **b)** Prevalence of co-colonization *I*_*ij*_ tends to scale with the product of strain prevalences in single colonization (*I*_*i*_, *I_j_*), for all strain pairs and for all time, when on the slow manifold (Table 1). This example is simulated using a random matrix *K*, with *N* = 10. Each trajectory corresponds to a given strain in the system (a), or a given strain pair (b). Another example for *N* = 20 is shown in Supplementary figure S7.

The fact that prevalence of co-colonization with strains *i* and *j* in this system, involves a product between single prevalences of *i* and *j* in the population (*I*_*ij*_ ~ *I*_*i*_*I*_*j*_), even though the strains are interacting, is contrary to the independence closure assumption under a purely statistical perspective, adopted heuristically in epidemic models (Kucharski et al., 2016). As we show here, depending on epidemiological details, the feedbacks between interacting strains may mathematically lead to overall multiplicative effects between individual and dual strain prevalences in colonization, with an explicit pre-factor determined by *R*_0_ and *k*.

To test the quality of the slow-fast approximation with respect to the original system, we verify that the error between the two is small. This is made precise via numerical simulations (Figure S4), where the neutral model is shown to be a good approximation of the original system in a fast time-scale, *o*(1/*ε*), and the slow-dynamics reduction a good approximation on the longer time-scale (*εt*). Finally, we show that this approximation is also advantageous in terms of efficient computation of dynamics for an arbitrary number of strains *N* (Figures S5-S6). While reinforcing the validity of our method, these two important features can aid parameter inference frameworks for epidemiological time-series in multi-strain systems, known to pose many challenges (Shrestha et al., 2011; Gjini et al., 2016).

Overall this multi-strain model, the slow-fast decomposition, and the key features outlined above provide a crucial missing link towards the full characterization of conservative multi-type SIS dynamics with interactions in co-colonization. Our conceptual and analytical framework suggests that in understanding multi-type communities, a closer integration between different temporal scales on one hand, and demographic vs. selective processes on the other, may be possible. Ultimately, the key mechanistic link with the replicator equation, uncovered here, could have wide-ranging applications from microbial ecology to complex systems and public health.

## Discussion

One of the fundamental questions in ecology and evolutionary biology is the generation and maintenance of biodiversity (Gause, 1934; MacArthur, 1967; Hubbell, 2001). Most theoretical approaches consider multi-species interactions with classical Lotka-Volterra ODE systems (Pascual et al., 2006; Mougi and Kondoh, 2012) or with evolutionary game theory models (Nowak and May, 1992; Traulsen and Nowak, 2006), where the interplay between cooperation and competition is key. Typically when more population structure is involved (e.g. in epidemiology (Kucharski et al., 2016)) and in high-dimensional spaces of diversity, analysis is very challenging, calling for new theoretical tools and methods, beyond numerical simulation. So far, neither Lotka-Volterra models, nor the replicator equation have been explicitly adopted in mathematical epidemiology, typically based on multi-type compartmental descriptions of host population structure, with only relatively recent attention on evolutionary dynamics (Day and Gandon, 2006) and typically focusing on virulence evolution (Berngruber et al., 2013).

In this study, we have bridged further such analytical fronts, by considering a new system of multi-type interactions that arise in SIS transmission dynamics with co-infection and altered susceptibilities. We have derived an explicit link with the replicator equation (Taylor and Jonker, 1978; Weibull, 1997; Hofbauer and Sigmund, 2003; Cressman and Tao, 2014) for strain frequencies, which in turn, is mathematically related to Lotka-Volterra multi-species models (Lotka, 1926; Volterra, 1926). The continuous replicator equation on *N* types is topologically equivalent to the Generalized Lotka-Volterra equation in *N* − 1 dimensions (Bomze, 1983, 1995; Hofbauer and Sigmund, 2003). Thus, by uncovering the replicator equation at the heart of our co-colonization system, we offer new and promising avenues of theoretical cross-fertilization betwen ecology, epidemiology and evolution.

We investigated how co-colonization interaction coefficients (*K*_*ij*_), be they cooperative or competitive on average (*k* above/below 1) and with arbitrary among-strain variation (*α*_*ij*_), drive coexistence in a system with *N* similar types. We obtained a timescale separation for the dynamics, which links explicitly variation in interaction traits among co-colonizing strains (*K*_*ij*_ = *k* + *εα*_*ij*_), with the slow timescale (*εt*) under which their coupled selective dynamics unfold. Model reduction for small deviations from neutrality allowed us to express strain frequencies on the slow manifold via only *N* equations (instead of *N* + *N*(*N* − 1)/2 equations in the full system). Strain frequencies can be easily re-mapped to the more complex epidemiological variables reflecting population structure.

### From pairwise invasion to collective coexistence

We have demonstrated that the dynamics of the full system can be expressed entirely in terms of pairwise invasion fitnesses of each of the strains (Eq. (8)). This is a novel and important finding that links mathematically pairwise outcomes to emergent community dynamics. Our results suggest that a bottom-up approach can be applied to understand and exactly predict community structure. In a recent experimental study, investigating assembly rules in microbial communities (Friedman et al., 2017), survival in three-species competitions was predicted by the pairwise outcomes with an accuracy of 90%. Yet, a similar level of accuracy in competitions between sets of seven or all eight species was harder to obtain, and required additional information regarding the outcomes of the three-species competitions. Although their ecological dynamics was based on the generalized Lotka-Volterra framework, and ours here on a multi-type SIS dynamics, the key to higher-dimensional dynamics may be in exploiting the full structure of the nonlinear coupling between pairwise invasion fitnesses, made explicit here.

### Environmental feedback from higher-order interactions

Higher-order interactions are expected to emerge whenever the presence of an additional species changes the interaction between two existing species, and can impact on the maintenance of diversity (Billick and Case, 1994). In our co-colonization system, modeling explicitly type interactions with two types of resources: susceptibles *S* and singly-colonized hosts *I*, a certain type of higher-order interactions arise naturally because of the indirect effects that altered susceptibilities between any pair *i* and *j* in co-colonization have on suppressing or augmenting the available resources *I*_*i*_ and *I*_*j*_ for the rest of the community, and thus when summed, contribute to mean fitness among everybody in the system. In this entangled network, the multiple types modulate their common environment through the changing term *Q* in (8), which can mean ‘deterioration’ of the environment if *Q* > 0 or ‘amelioration’ of the environment if *Q* < 0. This does not necessarily mean strains become more cooperative or competitive in epidemiological co-colonization, as the dynamics of *q* (in Eq. (5)) can be different from the dynamics of *Q*.

Our expression for strain frequency evolution ((8)) makes it also explicit that ‘environmental deterioration’ may be seen as a cost for the existing collective (since it reduces each strain’s rate of growth), but it serves as a protective mechanism against invasion by an outsider strain, and viceversa: ‘amelioration’ may on one hand seem like it benefits all strains, but on the other it also benefits any outsiders, which eventually may invade more easily. Central to these insights is having made explicit in this particular model the dependence on environmental dynamics of the selective dynamics between types, both in invasion fitness trait space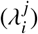, and in cocolonization trait space (*α*_*ij*_). This uncovers, in this system, the role of environmental feedback on eco-evolutionary processes, highlighted as a key challenge in evolutionary ecology (Lion, 2018).

### Invariant principles in N-type co-colonization

As known from frequency-dependent selection in evolutionary games, the final outcome among *N* players can be complex, represent a non-fitness-maximizing equilibrium and include oscillations and chaos (Nowak and Sigmund, 2004; Cressman and Tao, 2014). Yet, an important finding of our study concerns invariant principles emerging in non-equilibrium multi-type dynamics: the first one being about the dominance of types in single and co-colonization, which is expected to be equal, and the second one being about the co-colonization prevalences as a function of single colonization prevalences of strains (Figure 4, Supplementary Figure S6), and these could be used as a practical test for quasi-neutrality, when strain prevalence data are available. Our recapitulation of co-colonization dynamics from slow variables, sheds new analytical light on pathogen interactions and their manifestation at the epidemiological level (Kucharski et al., 2016), providing the link between within-host co-occurrence and population-level prevalences between strains. While independence underpins a majority of methods for detecting pathogen interactions from cross-sectional survey data (see e.g. (Valente et al., 2012)), it is being recognized that even simple epidemiological models challenge the underlying assumption of statistical independence (Hamelin et al., 2019). These studies are showing that even if pathogens do not interact, other epidemiological feedbacks can induce positive correlation between their prevalences, which leads the proportion of co-infected hosts to be higher than multiplication would suggest.

Along similar logic, our results here expose very clearly, that even if pathogens interact (e.g. via altered susceptibilities to coinfection), multiplicative effects between their prevalences emerge epidemiologically in co-colonization, but with an explicit pre-factor dependent on overall transmission intensity and mean interaction coefficient. Such finding invites a revision of methods to identify interactions between pathogens in endemic systems from cross-sectional data, based on a deeper mathematical understanding of underlying feedbacks. However, as already cautioned in previous studies (Cobey and Lipsitch, 2012b), patterns of interaction are subtle to detect from data and may require very large sample sizes to assess statistically, even with more sophisticated mathematical expectations.

### Extensions and outlook

In our system, microbial strains can infect a host simultaneously, and here we concentrated only on the case of up to 2 strains co-colonizing a host. Extension to higher multiplicity of infection (MOI) could be studied in the future, making use of previous models addressing arbitrary MOI and altered susceptibilities in co-infection (Adler and Brunet, 1991). The transmission and clearance rates of all strains here were assumed equal, except for the interactions in co-colonization given by an *N* × *N* matrix. In an ongoing work we show that this assumption can be relaxed leading to the *same* replicator equation with invasion fitnesses. The new invasion fitness expression emerging from these additional asymmetries in other traits is then a combination of all the deviations from neutrality (unpublished).

Past theoretical work has considered vulnerability to co-infection modeling it as a single mean-field parameter (Alizon, 2013). Other studies have studied how this trait at the host-pathogen interface impacts disease persistence (Gaivão et al., 2017), coexistence and vaccination effects (Lipsitch, 1997; Gjini et al., 2016), and contributes to diversity in other traits, e.g. virulence (Alizon et al., 2013) and antibiotic resistance (Davies et al., 2019). With the here-proposed analytical framework, exploration of these other biological aspects of systems with co-colonization could be made deeper and more insightful. We expect our results to stimulate further progress for the investigation of coexistence and evolution in multi-strain epidemiological systems such as the one of pneuomococcal bacteria (Cobey and Lipsitch, 2012a), where explicit mathematical results linking neutral and niche mechanisms, for high-dimensional coexistence, are still needed.

Our main goal here was to present the timescale decomposition for this system and highlight the fundamental features of the reduced model. While there is a lot of complex dynamics for different *N* and different matrix structures that we have not explored here, including multistability, limit cycles and chaos, our results thusfar clearly delineate promising avenues for new theoretical work. The wider and more complete ecological picture of co-colonization, as well as the gradients in diversity-stability regimes in coexistence is the focus of another study (Gjini and Madec, 2020). A natural next step is comparison with the classical Lotka-Volterra model (Lotka, 1926; Volterra, 1926), widely studied in ecology and adopted in applications (Mougi and Kondoh, 2012; Friedman et al., 2017), with increasing impact in the microbiome era. A great advantage however, of our framework, uncovering the replicator equation (Cressman and Tao, 2014) at the core of the dynamics, is because thanks to our mechanistic link, many mathematical results for general and special cases derived over decades for the replicator equation (and its links with LV models) (Hofbauer and Sigmund, 2003; Sandholm, 2010), would carry over automatically in this setting, and yield important insights for epidemiology.

The slow-fast framework allows to reduce the complexity of multi-type co-colonization systems, and understand transient and long-term coexistence on conserved manifolds near neutrality (Hubbell, 2001). Inevitably, in the quasi-neutral limit, delineated here, sampling noise is expected to become increasingly important. Utilisation of fast-variable elimination techniques is leading to growing awareness that noise-induced selection appears across several evolutionary systems in ecology, epidemiology and population genetics (reviewed in (Constable and McKane, 2018)). For instance, in a similar SIS model to ours, for two strains (Kogan et al., 2014), it was found that such noise generated selection for one of the strains, depending on the type of initial conditions, even though this was not predicted by the deterministic theory. Exploration of noise-induced selection during the fast (neutral) time-scale of our multi-strain system, where all strains are expected to behave as equivalent, would be interesting for the future.

In summary, although motivated by infectious disease transmission with altered susceptibilities in co-colonization (Gjini et al., 2016; Gjini and Madec, 2017), the global contagion dynamics captured here provide compelling parallels with other systems and should have broader applications. The coinfection model could be applied to study more mechanistically opinion propagation dynamics, coexistence in microbial consortia, plant ecology, and other multitype systems where colonizer-cocolonizer interactions matter. Thanks to its powerful abstraction and generality, this model and closed analytical form for frequency dynamics, offers a new bridge between population dynamics in epidemiology, community ecology, and Darwinian evolution.

## Supporting information

Supplementary material on mathematical derivations and additional figures

Movie of the dynamics in Figure 2.

Movie of more complex dynamics, possible in the system.

## APPENDIX 1: More details on Materials and Methods

Here, we briefly outline the materials and methods used. A more detailed treatment of the theoretical analysis and additional figures are presented in the online SI Appendix.

## Slow-fast representation of the dynamics

Since we model *N* closely-related strains, we can write each co-colonization coefficient as: *K*_*ij*_ = *k* + *εα*_*ij*_, where 0 ≤ *ε* ≪ 1. Replacing these in (3), and re-arranging, we obtain:

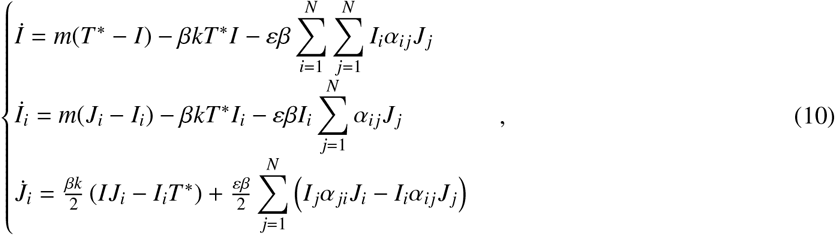

where 1 ≤ *i* ≤ *N*, and *I* = ∑ *I*_*j*_.

## Neutral system

If *ε* = 0, then we obtain the *Neutral model*, where all strain co-colonization coefficients are equal (*K*_*ij*_ ≡ *k*). The first equation above gives

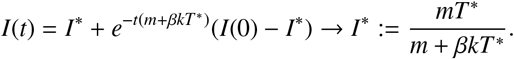

Co-colonization prevalence is simply derived as: *D*\* = *T*\* − *I*\*.

Fixing *I* = *I*\*, yields the linear system:

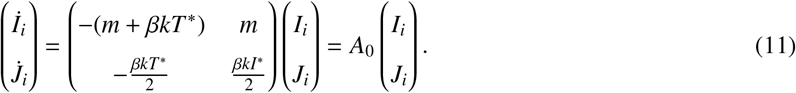

Matrix *A*_0_ has the two eigenvalues 0 and −*ξ* < 0. By defining *H*_*i*_ and *z*_*i*_ from the eigenvectors of *A*_0_ as:

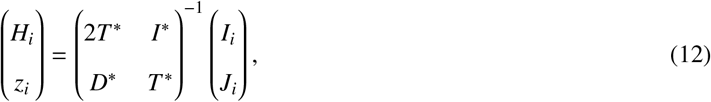

we have

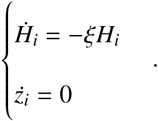

Thus on the fast time-scale

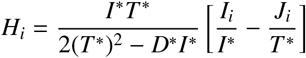

tends to zero, and

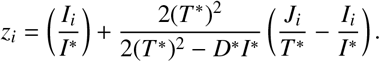

Because on the slow manifold, we have *H*_*i*_ = 0, we can infer that during fast dynamics *z*_*i*_ tends to:

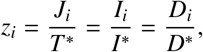

whereby *z*_*i*_ describes exactly the frequency of strain *i* in overall colonization, with ∑ *z*_*i*_ = 1.

## The slow dynamics: strain frequencies

Performing further analyses of system (10) for *ε* > 0, on the slow timescale *εt* (See Supplementary material 2), we find that *z*_*i*_ on 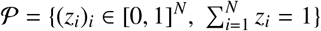, obey the explicit dynamics of (5):

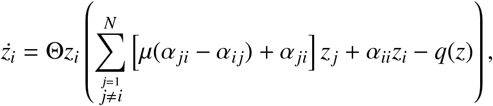

for 1 ≤ *i* ≤ *N*, where 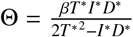; 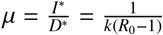, and the term *q*(*z*) is given by:

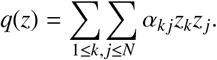

At this stage, we can apply quasi-stationarity methods Tikhonov (1952); Lobry and Sari (1998, 2005); Hoppensteadt (1966) to show that the solution of the full system tends to the solution of the slow-fast representation as *ε* → 0.

## Notes

#### Summary of Updates

We make explicit the link with the replicator equation. We elaborate on special key cases of mean fitness evolutionary dynamics. We highlight key insights and applications for microbial community analysis.

