## Supplementary material on mathematical derivations and additional figures for "Predicting *N*-strain coexistence from co-colonization interactions: epidemiology meets ecology and the replicator equation"

---

---

### 1. Table of illustrative model parameters for a biological system (e.g. pneumococcus bacteria transmission)

Table S1: Model parameters, some illustrative values, and their biological interpretation. Notice that the model is general enough to deal with any combination of parameters, provided they lead to a global endemic steady state and close values of co-colonization traits between strains.

| Parameter | Interpretation | Features | Illustrative value | Reference |
|---|---|---|---|---|
| $\beta$ | Transmission rate of pathogen | $\beta > 0$ | 2.3 $(\text{month})^{-1}$ | (Gjini et al., 2016) |
| $K_{ij}$ | Interaction coefficient between strains $i$ and $j$ in co-colonization | Altered susceptibility to secondary infection by $j$ when already colonized by $i$, relative to uncolonized host $\begin{cases} K_{ij} \in [0,1] \text{ competition} \\ K_{ij} > 1 \text{ facilitation} \end{cases}$ | $\approx 0.1$ | (Lipsitch et al., 2012) |
| $r$ | Susceptible recruitment rate | $r > 0$ | 0.02 $(\text{month})^{-1}$ | (Gjini et al., 2016) |
| $d$ | Natural host mortality rate | $d = r$ | 0.02 $(\text{month})^{-1}$ | |
| $\gamma$ | Clearance rate of colonization | $\gamma > 0$ | 0.8 $(\text{month})^{-1}$ | (Gjini et al., 2016) |
| $m$ | Net outflow rate from the colonization state | Inverse duration of any colonization episode | $m = r + \gamma$ | |
| $N$ | Number of strains in the system | Arbitrary, $N > 1$ | 30-60 | (Lipsitch et al., 2012) |
| $R_0$ | Basic reproduction number of the pathogen, $R_0 = \beta/m$, here assumed equal among strains | $\begin{cases} R_0 < 1 \text{ Disease extinction} \\ R_0 > 1 \text{ Persistence} \end{cases}$ | 2-3 | (Dietz, 1993),(Gjini et al., 2016) |

E. Gjini, C. Valente, R. Sá-Leão, and M. G. M. Gomes, Journal of Theoretical Biology **388**, 50 (2016).
M. Lipsitch, O. Abdullahi, A. D'Amour, W. Xie, D. M. Weinberger, E. T. Tchetgen, and J. A. G. Scott, Epidemiology **23**, 510 (2012).
K. Dietz, Statistical methods in medical research **2**, 23 (1993).

### 2. Derivation details for the slow-fast dynamics

Using the ansatz: $I(t) = \mathbf{I} + \varepsilon X(t) + O(\varepsilon)$, system (11) reads

$$\begin{cases} \dot{X} = -(m + \beta k\mathbf{T})X - \beta \sum_{i=1}^{n} \sum_{j=1}^{n} I_i \alpha_{ij} J_j + O(\varepsilon) \\ \dot{\begin{pmatrix} I_i \\ J_i \end{pmatrix}} = A \begin{pmatrix} I_i \\ J_i \end{pmatrix} + \frac{\varepsilon\beta k}{2} X J_i \begin{pmatrix} 0 \\ 1 \end{pmatrix} + \varepsilon\beta \mathbf{W}_i(\mathbf{I}, \mathbf{J}) + O(\varepsilon) \end{cases}$$

wherein we have denoted:

$$\mathbf{W}_i(\mathbf{I}, \mathbf{J}) = \begin{pmatrix} -\sum_j I_i \alpha_{ij} J_j \\ \frac{1}{2} \sum_j I_j \alpha_{ji} J_i - I_i \alpha_{ij} J_j \end{pmatrix}$$

The final slow fast form is given by using the new variables $H_i$ and $z_i$ defined in (13). Hence, define

$$\begin{pmatrix} h_{i,1}(\mathbf{I}, \mathbf{J}) \\ h_{i,2}(\mathbf{I}, \mathbf{J}) \end{pmatrix} = \begin{pmatrix} 2\mathbf{T} & \mathbf{I} \\ \mathbf{D} & \mathbf{T} \end{pmatrix}^{-1} \begin{pmatrix} -2 \sum_j \alpha_{ij} J_j \\ \sum_j I_j \alpha_{ji} J_i - I_i \alpha_{ij} J_j \end{pmatrix}$$

and let us introduce the $\mathbf{G}_i = (g_{i,1}, g_{i,2})^T$ [1] as:

$$g_{i,s}(\mathbf{H}, \mathbf{z}) = (2(\mathbf{T})^2 - \mathbf{ID}) h_{i,s}(2\mathbf{T}\mathbf{H} + \mathbf{I}\mathbf{z}, \mathbf{D}\mathbf{H} + \mathbf{T}\mathbf{z}), \qquad s = 1, 2.$$

and

$$Q(\mathbf{H}, \mathbf{z}) = \sum_{i,j} \alpha_{i,j} \left(2\mathbf{T} H_i + \mathbf{I} z_i\right) \left(\mathbf{D} H_j + \mathbf{T} z_j\right).$$

we have

$$\dot{X} = -(m + \beta k\mathbf{T})X - \beta Q(\mathbf{H}, \mathbf{z}) + O(\varepsilon)$$

Finally, using these new unknowns, one obtains the expression of the slow fast system (S1):

$$\begin{cases} \dot{X} = -(m + \beta k\mathbf{T})X - \beta Q(\mathbf{H}, \mathbf{z}) + O(\varepsilon) \\ \dot{H}_i = -\mu H_i + O(\varepsilon) \\ \dot{z}_i = \varepsilon\delta \left(g_{i,2}(\mathbf{H}, \mathbf{z}) + \mathbf{T}(\mathbf{D} H_i + \mathbf{T} z_i) X + O(\varepsilon)\right) \end{cases}, \quad 1 \le i \le N \tag{S1}$$

where $\delta = \frac{\beta}{2(\mathbf{T})^2 - \mathbf{ID}}$ is a positive constant. If $\varepsilon \to 0$ then $z_i$ remains constant and $(X(t), \mathbf{H}(t)) \to (\phi(\mathbf{z}), 0)$ where:

$$\phi(\mathbf{z}) = -\frac{\beta Q(0, \mathbf{z})}{m + \beta k\mathbf{T}} = -\frac{\beta \mathbf{IT}}{m + \beta k\mathbf{T}} \sum_{i,j} \alpha_{ij} z_i z_j. \tag{S2}$$

On the slow time scale $\tau = \varepsilon t$ in (S1) , and letting $\varepsilon \to 0$, one obtains the slow dynamics

$$\begin{cases} 0 = -(m + \beta k\mathbf{T})X - \beta Q(\mathbf{H}, \mathbf{z}) \\ 0 = -\mu H_i \\ \dot{z}_i = \delta \left(g_{i,2}(\mathbf{H}, \mathbf{z}) + k(\mathbf{T})^2 z_i X\right) \end{cases}, \quad 1 \le i \le N \tag{S3}$$

Taking $(X, \mathbf{H}) = (\phi(\mathbf{z}), 0)$ yields the slow dynamics reducted to the slow manifold

$$\dot{z}_i = \delta \left(g_{i,2}(0, \mathbf{z}) + \beta k\mathbf{T}\phi(\mathbf{z})\right), \quad 1 \le i \le N$$

[1] Denoting $\mathbf{G} = (\mathbf{G}_i)_{1 \le i \le N}$ and $\mathbf{W} = (\mathbf{w}_i)_{1 \le i \le N}$, this simply means that $\mathbf{G} = \phi^{-1} \circ \mathbf{W} \circ \phi$ where $\phi(\mathbf{H}, \mathbf{z}) = (\mathbf{I}, \mathbf{J})$ is the linear change of variables defined in (13).

where

$$\begin{aligned} g_{i,2}(0,\mathbf{z}) &= \mathbf{IT}\sum_{j=1}^{N}\left[\mathbf{D}z_i\alpha_{i,j}z_j + \mathbf{T}(z_j\alpha_{j,i}z_i - z_i\alpha_{i,j}z_j)\right] \\ &= \mathbf{ITD}\sum_{j=1}^{N}\left(\frac{\mathbf{T}}{\mathbf{D}}\alpha_{j,i} - \frac{\mathbf{I}}{\mathbf{D}}\alpha_{i,j}\right)z_i z_j \end{aligned} \tag{S4}$$

$\phi(\mathbf{z})$ is given by (S2) and verifies

$$k\mathbf{T}^2\phi(\mathbf{z}) = -\mathbf{IT}\frac{k\mathbf{T}^2}{m+\beta k\mathbf{T}}\sum_{i,j}\alpha_{ij}z_i z_j = -\mathbf{ITD}\sum_{i,j}\alpha_{ij}z_i z_j$$

Denoting $\mu = \mathbf{I}/\mathbf{D}$ and $\Theta = \dfrac{\beta\mathbf{TID}}{2(\mathbf{T})^2 - \mathbf{ID}}$, the explicit slow dynamics is then given by system (5) which may be rewritten as

$$\frac{d}{d\tau}\mathbf{z} = \Theta\mathbf{z}\cdot\left(\mu(A^t - A)\mathbf{z} + A^t\mathbf{z} - \mathbf{z}^t A\mathbf{z}\right). \tag{S5}$$

Remark that we have:

$$\frac{d}{d\tau}\sum_j z_j = \Theta\sum_{i,j}\alpha_{i,j}z_i z_j\left(1 - \sum_j z_j\right). \tag{S6}$$

which implies that the simplex $\mathcal{P} = \left\{\mathbf{z}\in\mathbb{R}_+^N,\ \sum z_j = 1\right\}$ is invariant under (5).

### 3. Characterization of the model via invasion fitnesses

We focus on the trivial solutions of (5), which are the extreme competitive exclusion equilibria $E_i = (\delta_{ik})_{1\le k\le n}$ where $\delta_{ik}$ is the Kronecker delta: $\delta_{ik} = \begin{cases}1 \text{ if } i = k,\\ 0 \text{ if } i \neq k.\end{cases}$

In particular, we note $\lambda_i^j$ the invasion fitness of the strain $i$ with respect to the trivial solution $E_j$. More precisely, $\lambda_i^j$ is the per-capita growth rate of the strain $i$ evaluated at $E_j$. Since the $i$-th line of the system reads $\frac{d}{d\tau}z_i = z_i h_i(z_1,\cdots,z_n)$, we have the explicit formula:

$$\lambda_i^j = h_i(E_j)$$

that is

$$\lambda_i^j = \mu(\alpha_{ji} - \alpha_{ij}) + \alpha_{ji} - \alpha_{jj}.$$

We have that the line $i$ of (S5) is

$$\dot{z}_i = z_i\left(\sum_{\substack{j=1\\ j\neq i}}^{N}\left(\mu(\alpha_{ji} - \alpha_{ij}) + \alpha_{ji}\right)z_j + \alpha_{ii}z_i - \sum_{j=1}^{N}\sum_{k=1}^{N}\alpha_{kj}z_k z_j\right).$$

With the notations introduced above, this can be rewritten as:

$$\dot{z}_i = z_i\left(\sum_{\substack{j=1\\ j\neq i}}^{N}\lambda_i^j z_j + \sum_{j=1}^{N}\alpha_{jj}z_j - \sum_{j=1}^{N}\sum_{k=1}^{N}\alpha_{kj}z_k z_j\right). \tag{S7}$$

From $\sum_{j=1}^{N} z_j = 1$, we infer

$$\begin{aligned}\sum_{j=1}^{n} \alpha_{jj} z_j - \sum_{k,j} \alpha_{kj} z_k z_j &= \sum_{j=1}^{N} z_j (\alpha_{jj} - \sum_{k=1}^{N} \alpha_{kj} z_k) \\ &= \sum_{j=1}^{N} z_j (\alpha_{jj} \sum_{k=1}^{N} z_k - \sum_{k=1}^{N} \alpha_{kj} z_k) \\ &= \sum_{j=1}^{N} \sum_{k=1}^{N} z_j (\alpha_{jj} - \alpha_{kj}) z_k\end{aligned}$$

finally, the fact that $\sum_{j=1}^{N} \sum_{k=1}^{N} (\alpha_{jk} - \alpha_{kj}) z_j z_k = 0$ yields

$$\sum_{j=1}^{n} \alpha_{jj} z_j - \sum_{k,j} \alpha_{kj} z_k z_j = - \sum \sum_{1 \le k \ne j \le N} \lambda_j^k z_j z_k.$$

Plugging this into (S7), we obtain a new version of system (S5), enterely parametrized by the fitnesses $\lambda_i^j$ :

$$\begin{cases} \frac{d}{d\tau} z_i = \Theta z_i \cdot \left( \sum_{j \ne i} \lambda_i^j z_j - \sum \sum_{1 \le k \ne j \le N} \lambda_j^k z_j z_k \right), \quad i = 1, \cdots, N \\ z_1 + \cdots + z_N = 1. \end{cases} \tag{S8}$$

or more shortly by noting $\lambda_i^i = 0$ and the *pairwise invasion fitness matrix* $\Lambda = (\lambda_i^j)_{1 \le i,j \le N}$ :

$$\begin{cases} \frac{d}{d\tau} \mathbf{z} = \Theta \mathbf{z} \cdot (\Lambda \mathbf{z} - \mathbf{z}^t \Lambda \mathbf{z}) \\ \\ z_1 + \cdots + z_N = 1. \end{cases} \tag{S9}$$

### 4. Special cases for mutual invasion fitnesses

*Symmetric invasion matrix*

In this case, starting with $\lambda_i^j = \lambda_j^i$, for all pairs of strains the following holds:

$$\frac{\alpha_{jj} - \alpha_{ii}}{\alpha_{ji} - \alpha_{ij}} = 2\mu + 1 > 1$$

This case resembles the classical condition for coexistence in ecology where the difference in within-strain interactions being bigger than that in between-strain interactions, favours coexistence between the strains, independently of whether the mean trait is cooperation or competition: the important principle is about deviation from the mean. If two strains cooperate more between than within-strain, then they will coexist. Similarly, if two strains compete more within strain than between strains, they will coexist.

In general, in this case, we have that $Q$ always increases: $\dot{Q} \ge 0$. Indeed, denote $\bar{\lambda}_i = \sum_j \lambda_i^j z_j$. We have

$$\dot{Q} = \sum_{i=1}^{n} \sum_{j \ne i} \lambda_i^j (z_i \dot{z}_j + z_j \dot{z}_i) = \sum_{i=1}^{n} \sum_{j \ne i} \lambda_i^j z_i z_j (\sum_{k \ne i} \lambda_i^k z_k + \sum_{k \ne j} \lambda_j^k z_k - 2Q)$$

Hence

$$\dot{Q} = (\sum_{i=1}^{n} z_i \sum_{j \ne i} \sum_{k \ne j} \lambda_i^j z_j \lambda_i^k z_k - Q^2) + (\sum_{j=1}^{n} z_j \sum_{i \ne j} \sum_{k \ne j} \lambda_i^j z_i \lambda_j^k z_k - Q^2)$$

We have $Q = \sum z_i \overline{\lambda}_i$ (i.e. mean invasibility trait in the system) and then

$$\sum_{i=1}^{n} z_i \sum_{j \neq i} \sum_{k \neq j} \lambda_i^j z_j \lambda_i^k z_k - Q^2) = var(\overline{\lambda}_i)$$

If moreover $\lambda^T = \Lambda$ it comes

$$(\sum_{j=1}^{n} z_j \sum_{i \neq j} \sum_{k \neq j} \lambda_i^j z_i \lambda_j^k z_k - Q^2) = (\sum_{j=1}^{n} z_j \sum_{i \neq j} \sum_{k \neq j} \lambda_j^i z_i \lambda_j^k z_k - Q^2) = var(\overline{\lambda}_i)$$

and then

$$\dot{Q} = 2var(\overline{\lambda}_i) \geq 0.$$

- *Special case of all $\lambda$s negative*
  The system will be characterized by multistability and competitive exclusion. $Q$ will be negative and will increase towards zero. (similar too the case of defense- driven invasion).
- *Will only the pairs with $\lambda_i^j = \lambda_j^i > 0$ coexist?* If $\lambda_i^j = \lambda_j^i < 0$ then the coexistence steady state $E_{i,j}$ exists but it is unstable. For a system with many strains, the outcomes can be complex.

*Invader-driven invasion matrix*

For this case, we find that there are many possible coexistence solutions subject to the constraint:

$$\lambda_i(1 - z_i) = \sum_{j=1}^{N} \lambda_j z_j (1 - z_j) = Q.$$

This leads to $z_i = 1 - \frac{Q}{\lambda_i}$ at steady state. Since $z_i < 1$ this yields that for all the strains $i$ which coexist, $Q$ and $\lambda_i$ have the same sign. So **the invasion fitnesses of the coexisting strains have the same sign**. If there is a steady state with a negative $Q$, and at least one member strain $j$ with positive invasion fitness, then the steady state is unstable for the growth rate of $j$ being in that case $\lambda_j - Q > 0$. This implies that for the steady state to be stable, either all the strains have negative fitness or the coexisting strains have positive fitness. In the first case, we have a multistability of competitive exclusion steady state and Q goes to 0. In the second case, then from $z_i > 0$, we have $\lambda_i > Q$ at equilibrium, and this means only the strains with a large enough intrinsic invasion fitness $\lambda_i$ will coexist.

In detail, at steady state with $n$ coexisting strains, one obtains $Q = \frac{n-1}{\sum_j \frac{1}{\lambda_j}}$, and thus for coexisting equilibrium strain frequencies:

$$z_i = 1 - \frac{n - 1}{\lambda_i \sum_j \frac{1}{\lambda_j}}.$$

*Resident-driven invasion matrix*

In this case the equation for $z_i$ can be written as:

$$\frac{dz_i}{d\tau} = \Theta z_i (\sum_{j \neq i} \lambda^j z_j - \sum_{j=1}^{N} \lambda^j z_j (1 - z_j)).$$

If a strain has negative invasion fitness, e.g. $\lambda^1 < 0$ (good defender of his equilibrium when alone), then the corresponding trivial steady state $E_1 = (1, 0, \cdots, 0)$ is asymptotically stable (because the linear growth rates of all other strains evaluated at such equilibrium, will always be negative. Hence, this situation leads often to a scenario of competitive exclusion with a single winner (and thus $Q(t) \to 0$). Several strains may have this property which leads to multistability of exclusion. Depending on initial conditions, there may exist some strain, for which $\sum_{j \neq i} \lambda^j z_j$ will be maximal compared to other strains in the system (the space of 'favourable' permissive territory is largest), and together with its initial frequency, this will drive the disproportionate growth of that strain in the system, leading to competitive exclusion, thus making $Q = 0$ in the long term. If the values of resident-driven fitnesses span the positive and negative range, and initial frequencies vary, different competitive exclusion scenarios may emerge, thus reinforcing the possibility of multi-stability.

However, there are exceptions. If a subset $I$ of $n$ strains have positive invasion fitnesses, they may coexist at a steady state $E_I$ and from $Q = \sum_{j \neq i} \lambda^j z_j$ and $\sum z_j = 1$ we infer that at such steady state:

$$z_i = \frac{1}{\lambda^i \sum_{j \in I} \frac{1}{\lambda^j}}$$

and $Q = \frac{n-1}{\sum_{j \in I} \frac{1}{\lambda^j}}$. But if this subset $I$ is strict ($n < N$), then this steady state $E_I$ is unstable because the other strains $i_0 \notin I$, at such equilibrium, will experience the positive growth rate

$$\sum_{j \in I} \lambda^j z_j - Q = \frac{1}{\sum_{j \in I} \frac{1}{\lambda^j}} > 0.$$

For example, if all the strains have a positive invasion fitness (everybody is a bad 'defender', thus everybody is invadable) then a stable coexistence of all $N$ types in the system is possible.

To conclude, for random $\lambda$ values which can be positive or negative, this type of matrix typically leads to a multistability of competitive exclusion steady states, although for very special cases, coexistence may arise.

*Anti-symmetric matrix*

Here we have for each couple of strains in the system $\lambda_i^j = -\lambda_j^i$. This leads to :

$$\frac{\alpha_{ii} + \alpha_{jj}}{\alpha_{ji} + \alpha_{ij}} = 1,$$

meaning that the cumulative (summed) deviation from the mean in within vs. between-strain interaction coefficients is equal for any two strains in the system, making the dynamics more difficult to settle, and thus complexity is more likely to arise in the multi-player community. For $Q$, we may write:

$$Q = \sum_i \sum_{j \neq i} (\lambda_i^j + \lambda_j^i) z_i z_j$$

In that case, it becomes immediately clear that $Q = 0$ at any time, independently on the values of type frequencies $z_i$.

### 5. Supplementary figures and movies of model dynamics

**Supplementary Movie S1. See separate file** *movie*1.*mp*4**.** Here we illustrate the dynamic trajectory of the system (Figure 2 of the paper). A. Network of pairwise invasion fitnesses between strains, where edges can be of 4 types: coexistence (red), bistability (blue), exclusion of $i$, exclusion of $j$ (directed gray arrows toward the winner). We plot the trajectory by considering the $z = (z_1, \cdots, z_6)$ as the barycentric coordinate of a point $M$ into the regular polygon determined by the nodes of the graph. B. Trajectory of the strain frequencies over time.

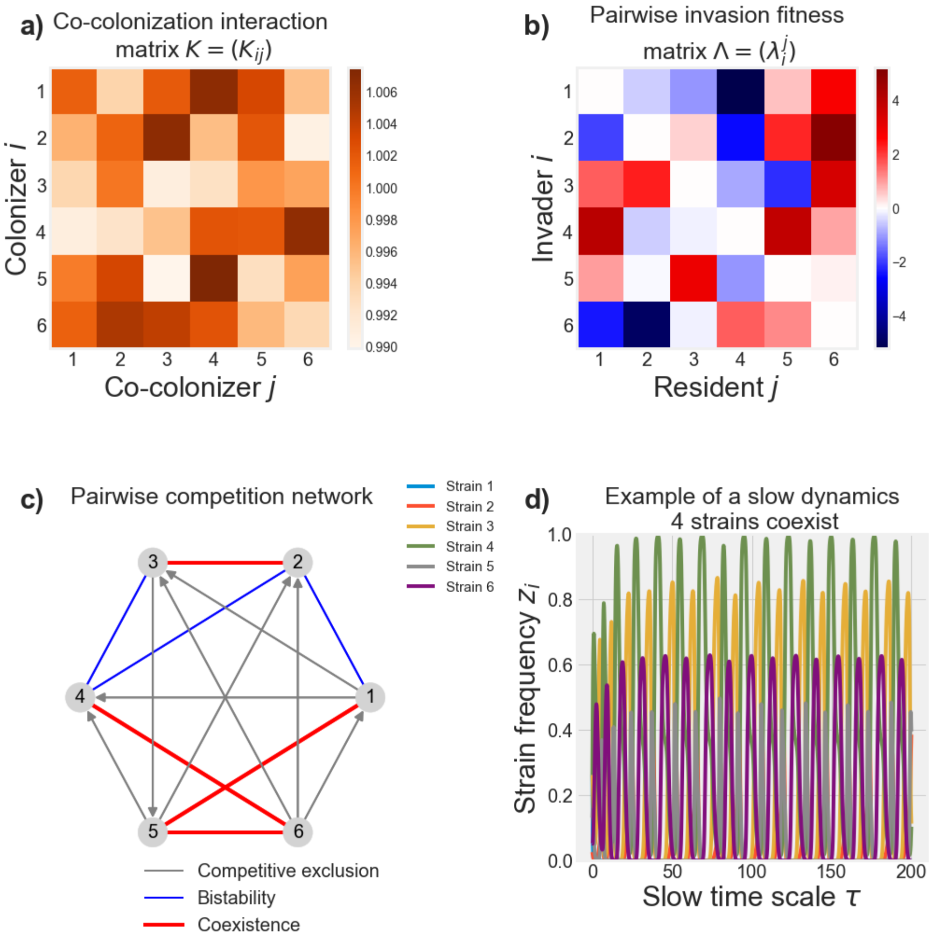


Figure S1: **Illustration of more complex dynamics**. Here we simulate the model in its reduced form for another set of parameters, leading to limit cycle coexistence between 5 strains in an $N = 6$ system. See associated movie S2. The precise co-colonization interaction matrix is the following:

$$K = \begin{pmatrix} 1.07961369 & 1.07350833 & 1.08150614 & 0.88653347 & 1.06682671 & 0.96613134 \\ 1.02917193 & 1.00960928 & 0.8335129 & 0.94718475 & 1.03909995 & 1.0328224 \\ 0.98795102 & 1.22187406 & 1.0750151 & 0.97995778 & 0.92834631 & 1.02456781 \\ 0.94758454 & 1.20872406 & 0.93043242 & 1.12848692 & 1.08616397 & 0.96457127 \\ 0.9974808 & 1.0684712 & 1.08323524 & 1.14132243 & 0.93105662 & 0.96004077 \\ 1.05293646 & 0.99303792 & 1.12707742 & 0.92449822 & 0.98562283 & 1.00990556 \end{pmatrix}.$$

**Supplementary Movie S2 (See separate file** *movie2.mp4***)** Here we illustrate the dynamic trajectory of the system (related to Figure S1). A. Network of pairwise invasion fitnesses between strains, where edges can be of 4 types: coexistence (red), bistability (blue), exclusion of $i$, exclusion of $j$ (directed gray arrows toward the winner). The system trajectory $z_i$ is shown as a movement in a 6-dimensional space. B. Trajectory of the strain frequencies over time.

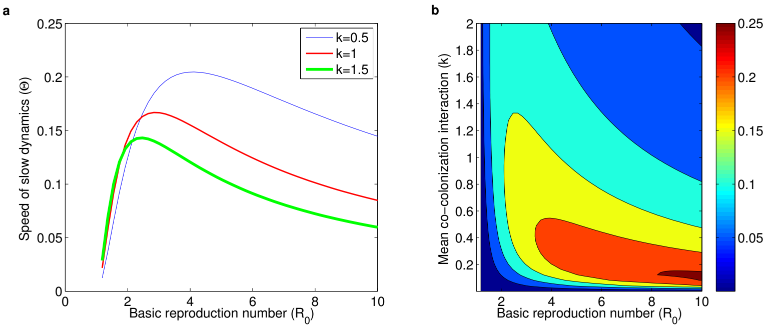


Figure S2: **Speed of slow dynamics depends on $R_0$ and k. a)** Values of $\Theta$ as a function of basic reproduction number $R_0$ for three values of mean interaction trait between strains in cocolonization $k$. The constant $\Theta$ is the rate that sets the tempo of multi-type "motion" on the slow manifold towards an equilibrium. $\Theta$ depends specifically on the absolute transmission rate of the pathogen $\beta$, which calibrates the slow dynamics, but also on mean traits $R_0$ and $k$, via the conserved aggregated quantities **T**, **I**, **D**, where $R_0$ and $k$ appear in nonlinear coupling with each other. There is an intermediate value of transmission intensity that maximizes the speed of strain frequency dynamics, and this increases with more competition between strains ($k \to 0$). **b)** $\Theta$ as a function of both $R_0$ and $k$. The speed of slow dynamics is maximal for intermediate values of $R_0$ and relatively low values of $k$. It is when competition dominates in co-colonization processes, that prevalence of singly-colonized hosts increases most in the system, speeding up the competitive dynamics between strains in the longer time scale. Here $\beta = 2$.

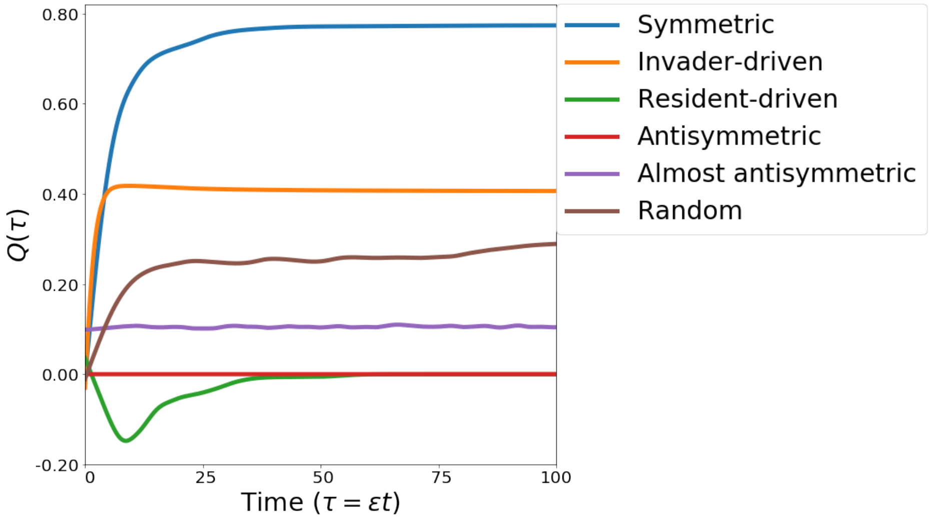


Figure S3: **Evolution of mean fitness $Q$ depends on pairwise invasion matrix structure.** This figure shows the mean behavior of the curves in Figure 3 of the main text. The fastest increase in mean fitness occurs for symmetric pairwise invasion fitnesses, whereas the slowest increase for random invasion. Mean fitness $Q$ indicates community resistance to invasion by outsider strains, and thus the system is more resistant when $Q$ is higher.

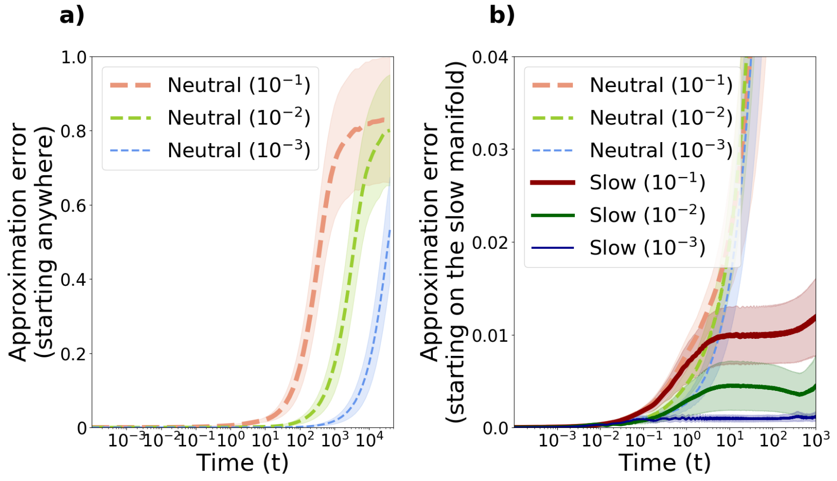


Figure S4: **Accuracy of the slow-fast approximation over time** We confirm there exists $t_0 > 0$ such that for any $T > t_0$ and any $\varepsilon > 0$ small enough, there is $M > 0$ such that for any $\tau \in (t_0, T)$, the error between the two is small: $\|S\left(\frac{\tau}{\varepsilon}\right) - \mathbf{S}\| + \sum_{i=1}^{N} \|I_i\left(\frac{\tau}{\varepsilon}\right) - \mathbf{I} z_i(\tau)\| + \sum_{1\leq i,j\leq N} \|I_{ij}\left(\frac{\tau}{\varepsilon}\right) - \frac{\mathbf{IT}}{\mathbf{S}} z_i(\tau) z_j(\tau)\| \leq M\varepsilon$. **a)** Error between the neutral model and the full system, starting from random initial conditions. We summarize simulations for different $\varepsilon$, where for each $\varepsilon$, 20 co-colonization matrices of different size ($N \in [2, 15]$) were randomly generated, together with initial conditions. On a fast time-scale ($o(1/\varepsilon)$) the neutral model is a very good approximation to the real system. **b)** Error of the neutral and the slow dynamics approximation, starting from the slow manifold (4). The error (mean over all model variables) between the original system trajectories and the neutral vs. slow dynamics. We generated random co-colonization matrices ($N \in [2, 15]$) for 3 values of $\varepsilon$ ($10^{-1}$,$10^{-2}$,$10^{-3}$) while keeping mean $k = 1$. The $K_{ij}$ were drawn from the normal distribution $N(k, \varepsilon^2)$. On the slow manifold, the slow-dynamics approximate well the original system, unlike the neutral model.

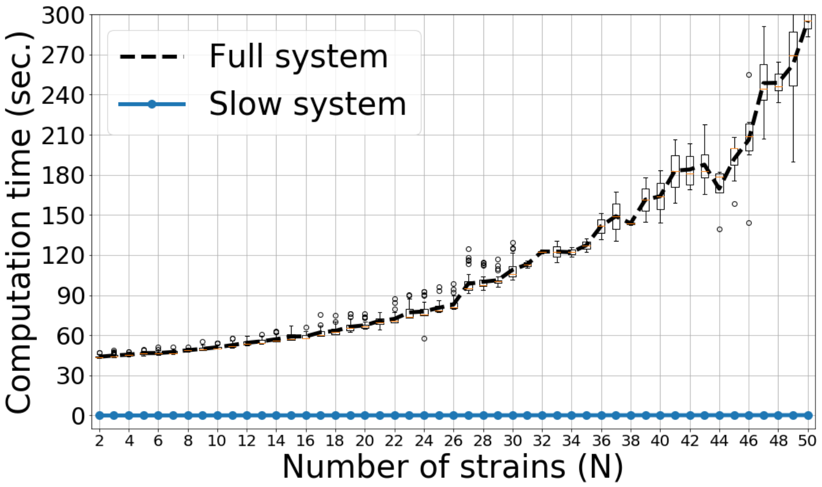


Figure S5: **Computation speed of the approximation.** Here, we verify that the slow-fast approximation enables exploration of dynamics in a much more computationally efficient manner over a very large number of strains $N$, which is difficult with the original system. Computation time increases as a function of number of strains $N$ in the full system (black dashed line) but not in the slow-fast dynamic aproximation (blue solid line) for $T = 5000$ time units. The line shows the mean over 20 random matrices for each $N$, with $\varepsilon = 0.1$, $k = 1$ and $R_0 = 2$. For details see Figure S6.

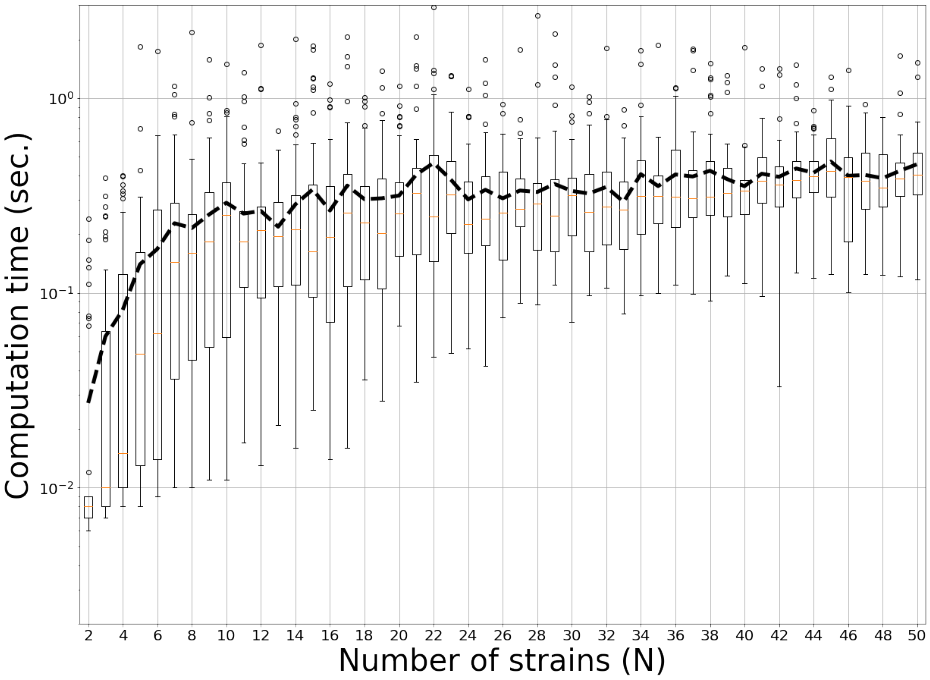


Figure S6: **Computation speed of the slow-dynamics approximation in detail.** Here we plot the computation time as a function of number of strains $N$, in terms of number of seconds on a PC to simulate the slow-fast dynamic aproximation (boxplots) for $T = 5000$ time units starting from the slow manifold. The line shows the mean over 20 random matrices for each $N$, where $\varepsilon = 0.1$ and mean co-colonization coefficient $k = 1$ and $R_0 = 2$.

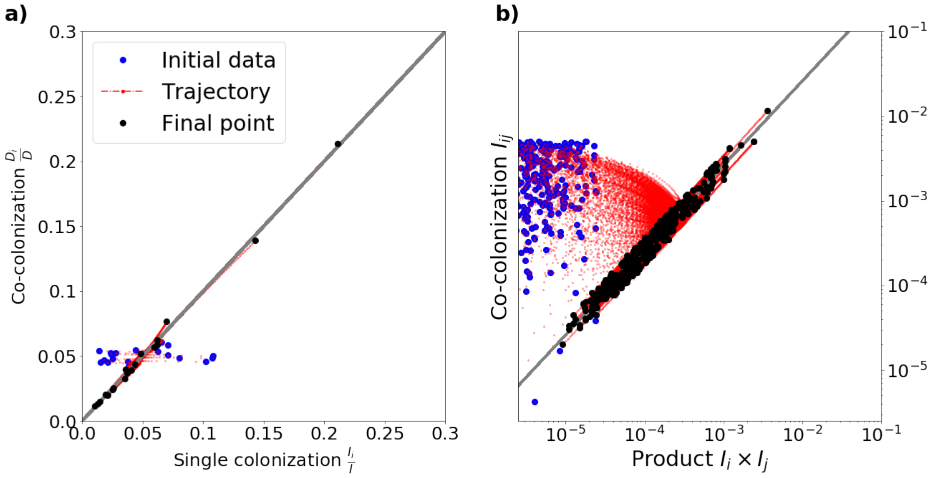


Figure S7: **Illustration of quasi-neutrality principles on the slow manifold for $N = 20$. a)** Strain frequencies tend to be the same in single and co-colonization, for all strains and for all time when on the slow manifold. **b)** Prevalence of co-colonization $I_{ij}$ tends to scale with the product of strain prevalences in single colonization ($I_i$, $I_j$), for all strain pairs and for all time, when on the slow manifold. This example is simulated using a random matrix $K$, with $N = 20$. Each trajectory corresponds to a given strain in the system (a), or a given strain pair (b).
